## Supplementary Figures & Methods for "Sphingosine 1-Phosphate Mediates Adiponectin Receptor Signaling Essential For Lipid Homeostasis And Embryogenesis"

RUNNING TITTLE: S1P promotes membrane homeostasis

AUTHORS: Mario Ruiz[1*], Ranjan Devkota[1], Dimitra Panagaki[1], Per-Olof Bergh[2], Delaney Kaper[1], Marcus Henricsson[2], Ali Nik[1], Kasparas Petkevicius[3], Johanna L. Höög[1], Mohammad Bohlooly-Y[3], Peter Carlsson[1], Jan Borén[2], Marc Pilon[1*]

AFFILIATIONS:
[1]Dept.Chemistry and Molecular Biology, Univ. Gothenburg, 405 30 Gothenburg, Sweden,
[2]Dept. Molecular and Clinical Medicine/Wallenberg Laboratory, Institute of Medicine, Univ. of Gothenburg, 414 67 Gothenburg, Sweden,
[3]Discovery Biology, Discovery Sciences, R&D, AstraZeneca, Gothenburg, Sweden.

CORRESPONDING TO: * Mario Ruiz or Marc Pilon, Dept. Chem Mol Biol, Univ. Gothenburg, Box 462, Gothenburg, SE-405 30, Sweden. Tel: +46 31 786 4952.

**Abstract**

Cells and organisms require proper membrane composition to function and develop. Phospholipids are the major component of membranes and are primarily acquired through the diet. Given great variability in diet composition, cells must be able to deploy mechanisms that correct deviations from optimal membrane composition and properties. Here, using lipidomics and unbiased proteomics, we found that the embryonic lethality in mice lacking the fluidity regulators Adiponectin Receptors 1 and 2 (AdipoR1/2) is associated with aberrant high saturation of the membrane phospholipids. Using mouse embryonic fibroblasts (MEFs) derived from AdipoR1/2-KO embryos, human cell lines and the model organism *C. elegans* we found that, mechanistically, AdipoR1/2-derived sphingosine 1-phosphate (S1P) signals in parallel through S1PR3-SREBP1 and PPARγ to sustain the expression of the fatty acid desaturase SCD and maintain membrane properties. Thus, our work identifies an evolutionary conserved pathway by which cells and organism achieve membrane homeostasis and adapt to a variable environment.

**Supplementary Information**

- Supplementary Fig. 1-8
- Supplementary methods (qPCR primer list and *C. elegans* methods)
- Supplementary References

Supplementary Fig.1 (related to Fig.1).

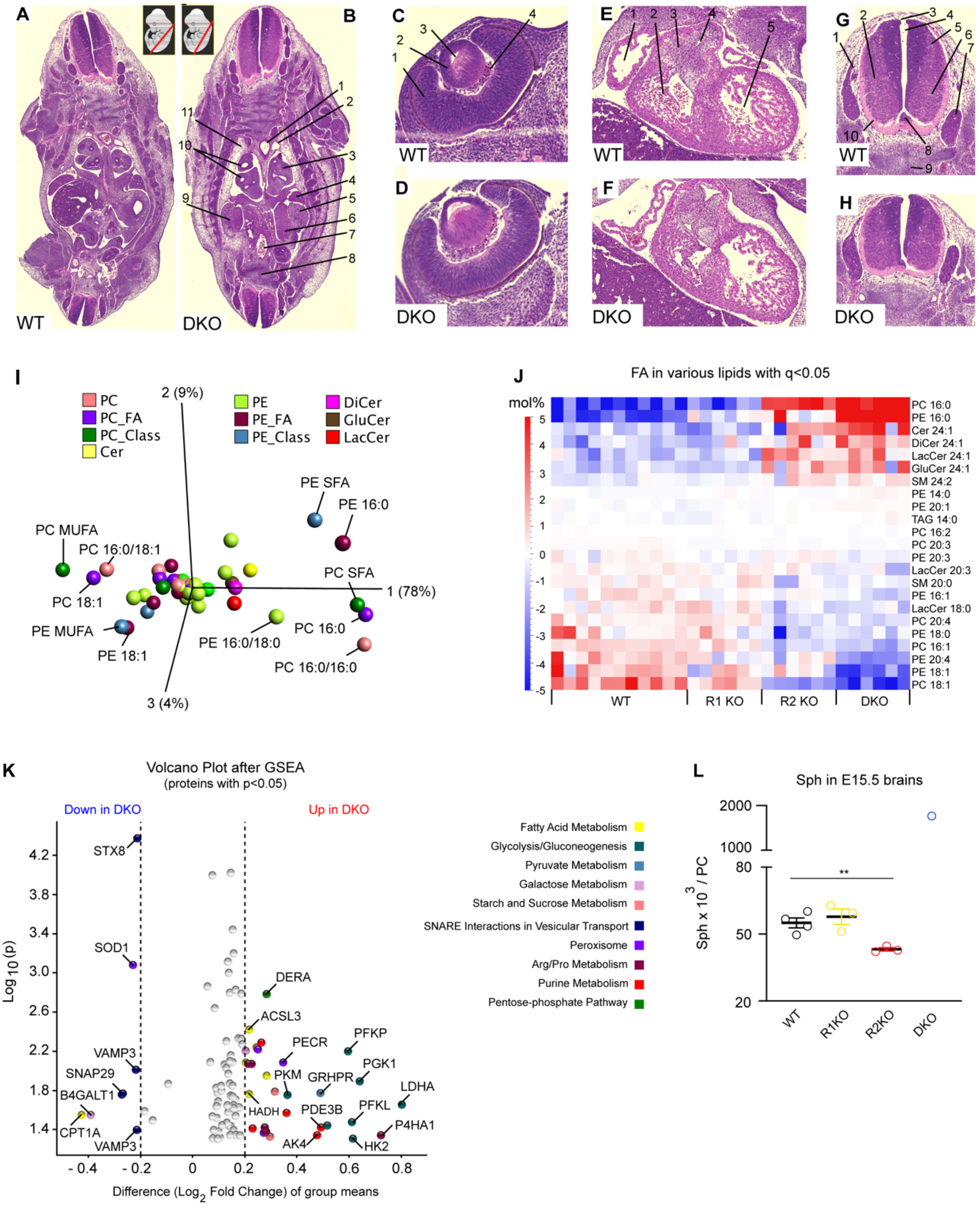

**Supplementary Fig.1 Membrane Lipid Composition Defects Precede Embryonic Lethality in DKO Mice** (related to Fig.1).

**(A-H)** Sections of WT and DKO embryos at E12.5 stained with H&E. In C: 1. thoracic (descending) aorta; 2. left subcardinal vein; 3. segmental bronchus (left lung); 4. left lobe of liver; 5. left metanephros (kidney); 6. rete ovarii (early stage of differentiation); 7. sacral vein; 8. notochord; 9. right metanephros (kidney); 10. segmental bronchus (right lung); 11. loose connective tissue forming wall of pericardio-peritoneal cavity. In D: 1. inner (neural) layer of retina; 2. lens; 3. lens vesicle; 4. hyaloid cavity. In F: 1. right atrium; 2. right ventricle; 3. leaflets of pulmonary valve; 4. origin of pulmonary trunk; 5. left ventricle. In H: 1. dorsal nerve root; 2. mantle; 3. roof plate; 4. central canal; 5. alar plate; 6. basal plate; 7. dorsal root ganglion; 8. floor plate; 9. notochord; 10. marginal zone.

**(I)** PCA showing the variables responsible for the genotype separation in Fig.1D.

**(J)** Heatmap of all the lipid species with a $q<0.05$ when comparing WT, R1KO, R2KO, DKO embryos at E12.5. Variables considered as in panel D. Note that WT embryos were enriched in PC with 16:1 and PE with 18:1 (both MUFA) whereas DKO had elevated PC with 16:0 and PE with 16:0 (SFA). Related lipidomics are included in Supplementary Data 2.

**(K)** Volcano plot of the proteomics showing all significant proteins after GSEA (97 in total), between WT and DKO at E12.5. Note that only the proteins with a log2 difference > than 0.2 from WT and with a $p<0.05$ are colored and labeled. See Supplementary Data 3 for the full list.

**(L)** Sph abundance in mouse embryonal brains at E15.5. Related lipidomics are shown Supplementary Data S2.

See also Supplementary Data 1-3. Data are represented as mean ± SEM. **$p<0.01$.

Supplementary Fig.2 (related to Fig.2).

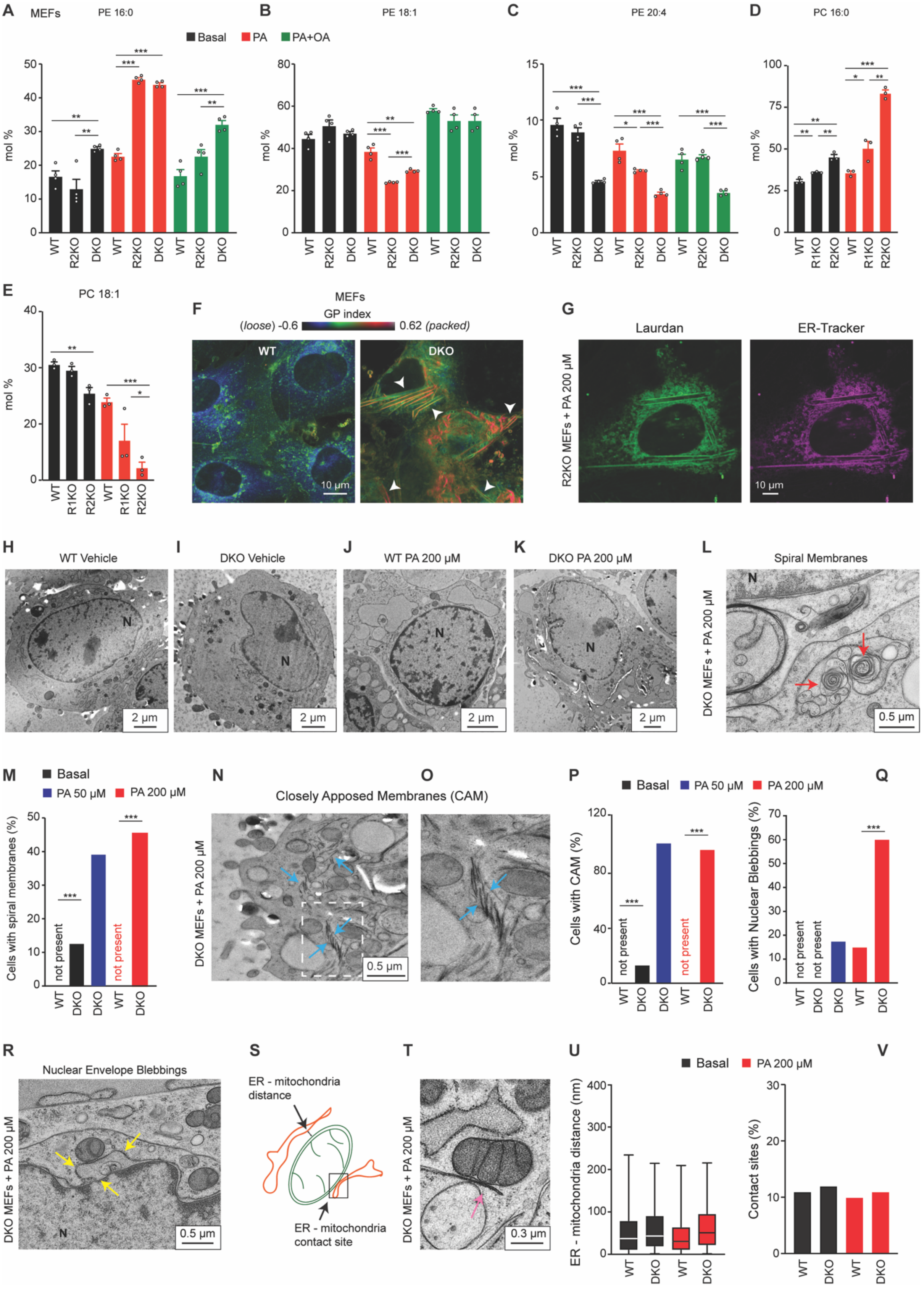

**Supplementary Fig.2 S1P Rescues Membrane Homeostasis Defects Caused by the Absence of AdipoR2 in MEFs** (related to Fig.2).
**(A-E)** PA (16:0), OA (18:1) and AA (20:4) abundance (mol%) in the PC and/or PE of WT, R1KO R2KO and DKO MEFs cultivated in the presence of vehicle, PA 200 µM ± OA 200 µM for 18 h. Related lipidomics are shown in Supplementary Data 4.
**(F)** Pseudocolor images of WT and DKO MEFs challenged with PA 200 µM and stained with Laurdan. Note that pixels with highly packed lipids are colored in red and appear only in DKO MEFs. White arrows indicate long and packed structures exclusively present in DKO cells. **(G)** Single channel confocal images of live R2KO MEFs stained with Laurdan (left picture) and ER-Tracker (right picture). Merged image is shown in Fig.2I.
**(H-K)** Electron microscopy pictures with a general cellular view of WT and DKO MEFs exposed to vehicle (F-G) and PA 200 µM (H-I).
**(L-M)** DKO MEFs images showing distinct spiral membranes (marked by red arrows) in the cytoplasm and the quantification (n=22-27 sections). N in the pictures indicates the nucleus.
**(N-P)** Representative image (and zoom in) of DKO MEFs exposed to PA 200 µM and showing electron dense closely apposed membranes (CAM, labeled with blue arrows) and the quantification (n=22-27 sections). The white square in N indicates the region magnified in O.
**(Q-R)** Representative image of DKO MEFs treated with PA 200 µM and showing nuclear envelop blebbing (yellow arrows) and the quantification (n=17-27 sections). N in the pictures indicates the nuclei.
**(S-V)** Cartoon, representative EM image and quantification of ER-mitochondria distance (n≈100) and contact-sites in WT or DKO MEFs in basal media and treated with PA 200 µM. Data are represented as mean ± SEM. *$p<0.05$, **$p<0.01$, ***$p<0.001$. See also Supplementary Data 4.

Supplementary Fig.3 (related to Fig.2).

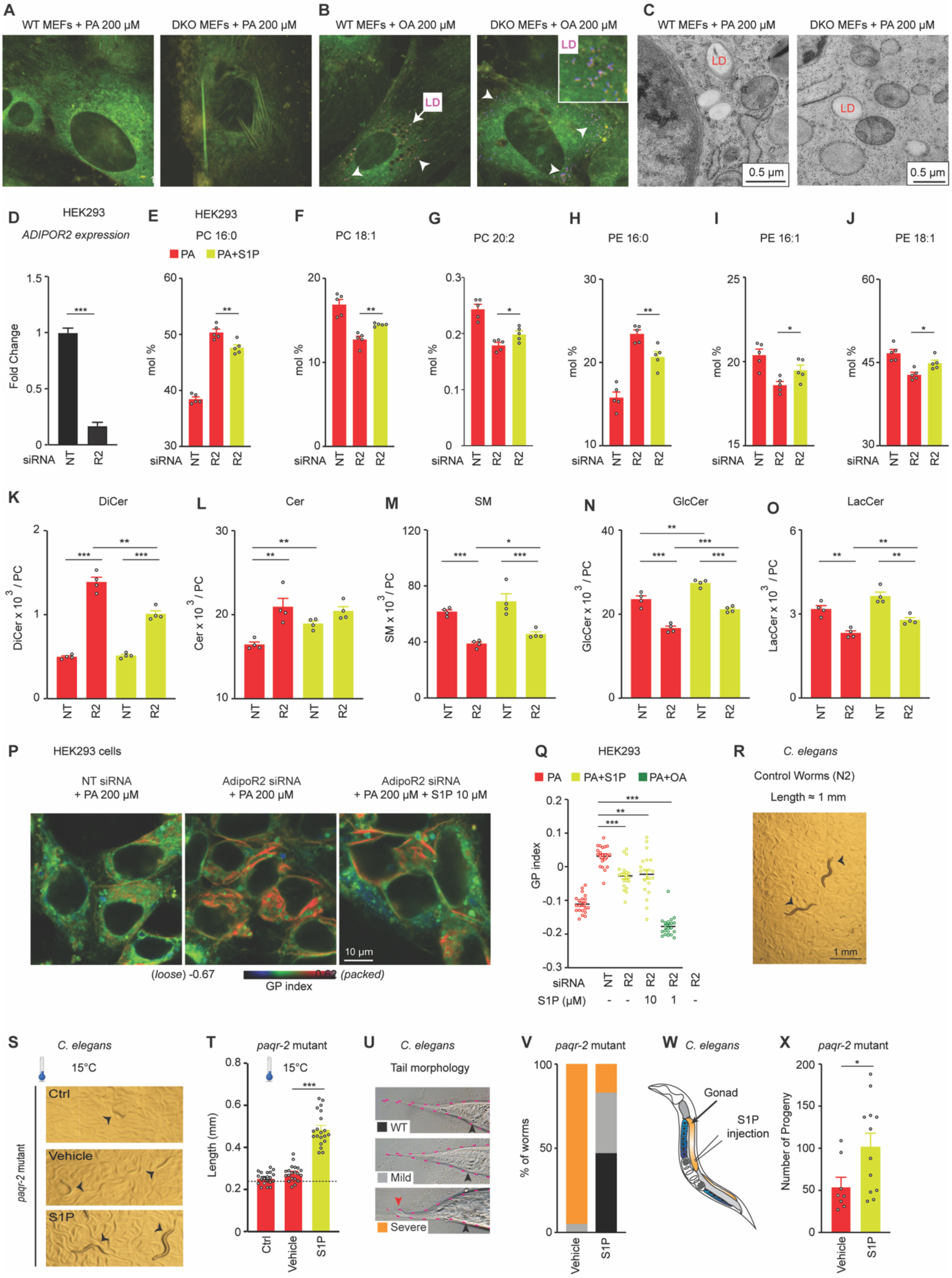

**Supplementary Fig.3 Loss of AdipoRs Causes Multiple Membrane Defects in MEFs, HEK293 Cells and in *C. elegans*** (related to Fig.2)**.**

**(A-B)** Representative images of MEFs stained with Laurdan (green and red color) and LipidSpot 610 (purple dots pointed by white arrows).

**(C)** Representative EM images of MEFs.

**(D)** qPCR showing the efficiency of the AdipoR2 knockdown in HEK293 cells.

**(E-J)** PA 16:0), OA (18:1) and eicosadienoic (20:2n-6) fatty acid abundance (mol%) in the PC and PA, PalOA (16:1) and OA in the PE of HEK293 cells cultivated in the presence of PA 200 µM ± S1P 1 µM and treated with NT and AdipoR2 siRNA. Related Lipidomics are shown in Supplementary Data 6.

**(K-O)** DiCer, Cer, SM, GlcCer and LacCer in HEK293 cells cultivated in the presence of PA 200 µM ± S1P 1 µM and treated with NT or AdipoR2 siRNA. From Supplementary Data 6.

**(P-Q)** Pseudocolor images and GP index of HEK293 cells treated with PA 200 µM ± S1P (10 and 1 µM) or ± OA 200 µM and stained with Laurdan dye.

**(R)** Representative picture of WT 1-day adult *C. elegans*.

**(S-T)** Representative pictures and quantification of the length of *paqr*-2 mutant worms (arrowheads) grown on control plates or vehicle ± S1P 25 µM at 15°C for 144 h. The dashed line in S represents the approximate length of the L1s at the start of the experiments.

**(U-V)** Representative images and quantification (n=50 worms/genotype) of the tail tip morphology of 1-day adult *C. elegans*: WT phenotype, and mild and severe defects are shown. Shorter and broader tail tips were scored as mild defect, whereas deformed and bumpy tails were scored as severe defect (indicated by the red arrow). Black arrows indicate the anus.

(**W-X**) Schematic representation of *C. elegans*. The region of the gonad where S1P was injected is highlighted in orange. Brood size of *paqr-2* mutant injected with vehicle ± S1P 10 µM.

Data are means ± SEM, except in D: ± SD. $*p<0.05$, $**p<0.01$, $***p<0.001$. See also Supplementary Data 6.

Supplementary Fig.4 (related to Fig.3).

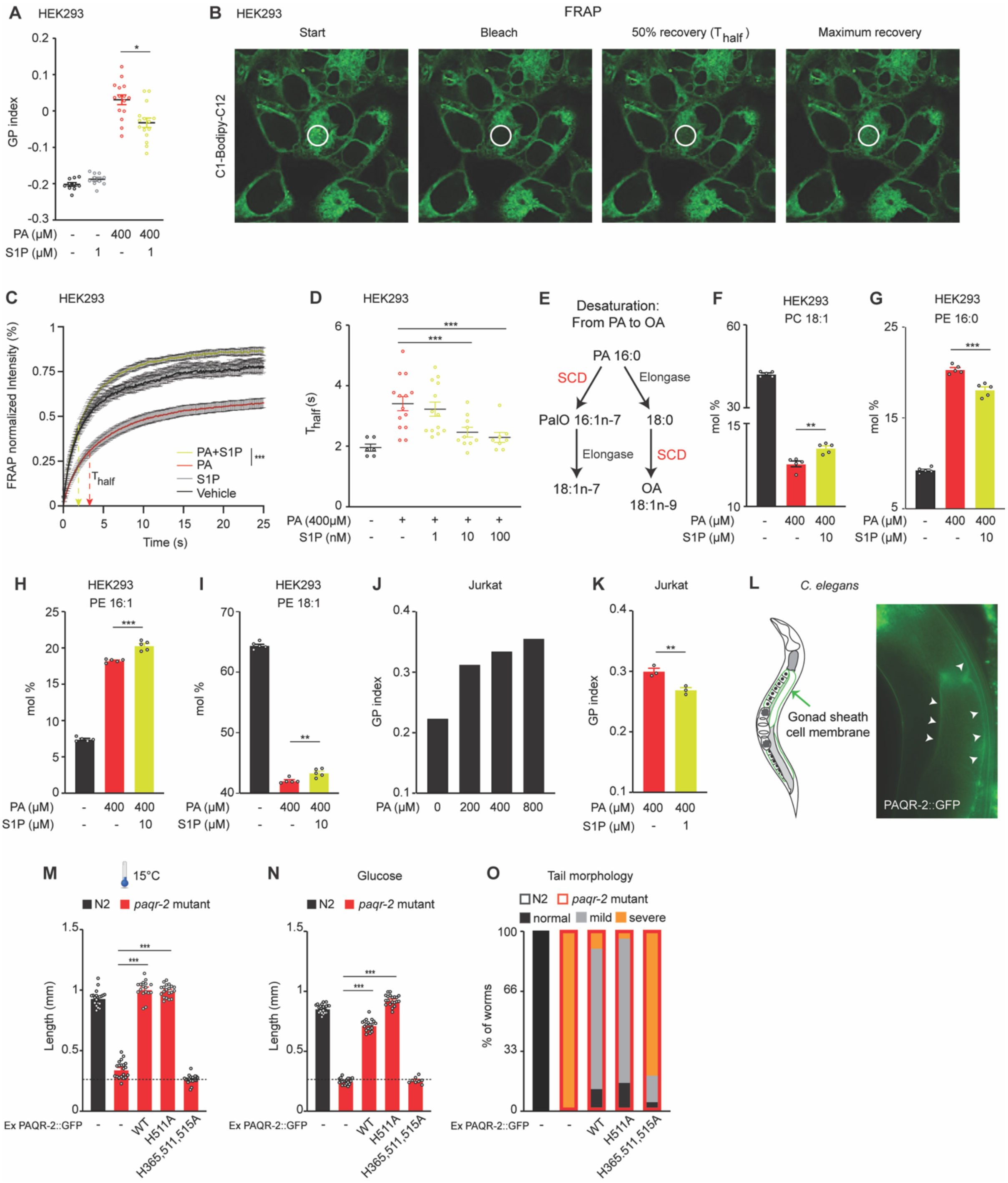

**Supplementary Fig.4. S1P Promotes Membrane Homeostasis in Multiple Human Cell Types and in *C. elegans*** (related to Fig.3).

**(A)** Average GP index from several images of HEK293 cells treated as indicated.

(**B-D)** C1-Bodipy-C12 staining of live HEK293 cells indicating different phases during a FRAP experiment (start, bleach, 50% recovery and end). The white circle represents the FRAP region. Panel C shows the normalized fluorescence intensity during the FRAP experiment of cells treated with vehicle, S1P 1 μM, and PA 400 μM ± S1P 1 μM. Panel D shows the average Thalf values from the curves in C.

**(E)** Pathway showing how PA (16:0) is desaturated (and elongated) to become 18:1.

**(F-I)** PalO (16:1) and OA (18:1) abundance (mol%) in the PC and PE of HEK293 cells treated as indicated. Data from Supplementary Data S5.

**(J-K)** Average GP index from cytometry experiments of Jurkat E6.1 cells treated with PA (panel J) or with PA 400 μM ± S1P (panel K) for 18 h and stained with Laurdan.

**(L)** Representation of *C. elegans* highlighting the gonad sheath cell membrane in green (tissue were PAQR-2::GFP is easily visualized) and a fluorescent picture of the actual gonad sheath cell membrane (white arrows).

**(M-N)** Length of N2 and *paqr-2* mutant worms expressing PAQR-2::GFP with WT sequence or with H511A and H365,511,516A variants. Worms were grown at 15°C for 144 h (panel M) or at 20°C on glucose plates for 72 h (panel N). Note that glucose is readily converted to SFAs by the dietary *E. coli*(Devkota et al., 2017). The dashed line represents the approximate length of the L1s at the start of the experiments.

**(O)** Quantification of the tail tip morphology of 1-day adult N2 and *paqr-2* mutant worms expressing different PAQR-2::GFP variants (n=50 worms/genotype DK). Examples of the different tail morphologies are shown in Supplementary Fig.3U.

**(P-Q)** qPCR results showing the efficiencies of the Sphk1 and Sphk2 knockdowns by siRNA in HEK293 cells.

Data shows mean ± SEM. *$p<0.05$, **$p<0.01$, ***$p<0.001$. See also Supplementary Data 5-S6.

Supplementary Fig.5 (related to Fig.4).

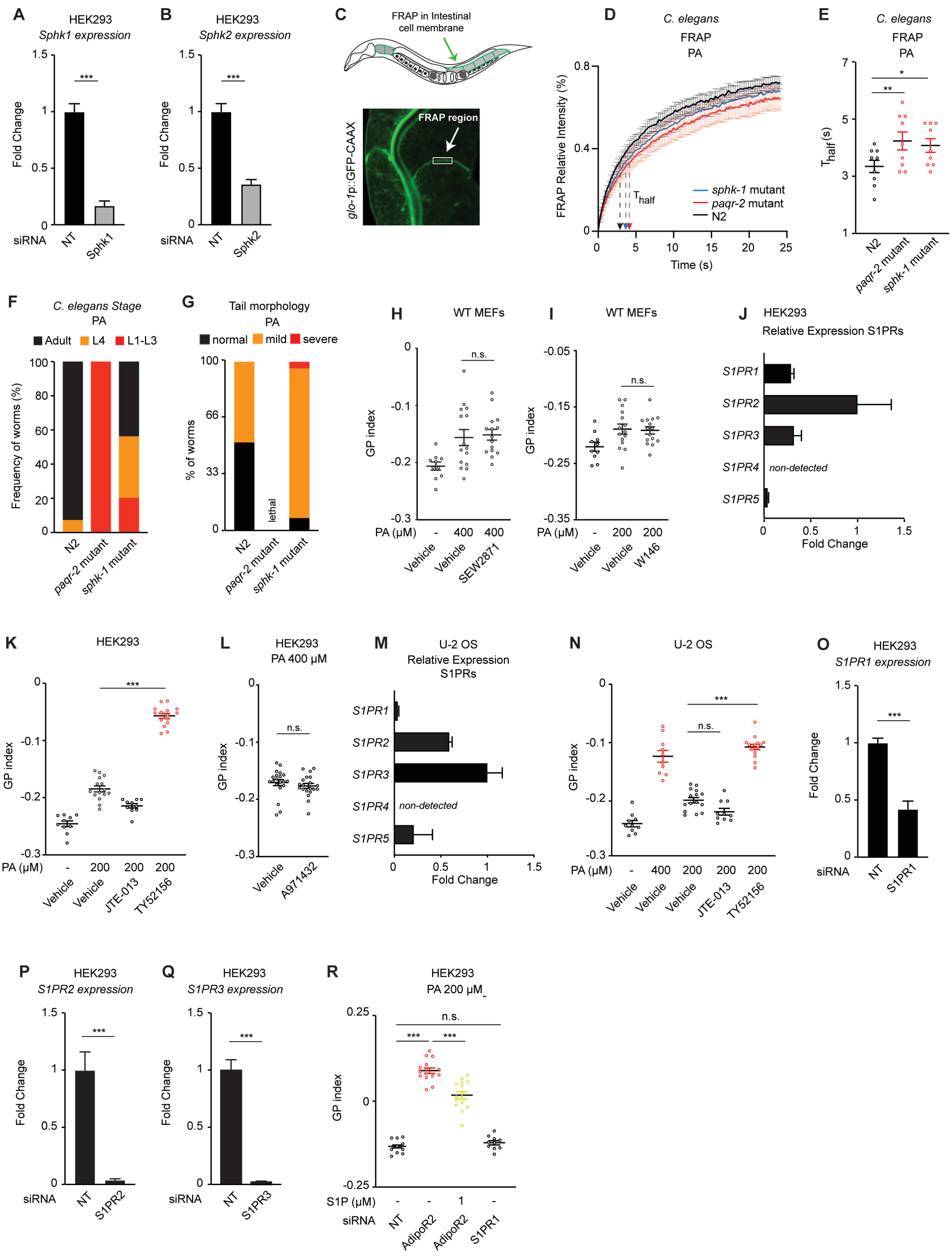

**Supplementary Fig.5 Sphingosine Kinases and S1PR3 Are Required to Maintain Membrane Homeostasis in Mammalian Cells and *C. elegans* (**related to Fig.4)

**(A-B)** qPCR showing the efficiencies of the Sphk1 and Sphk2 knockdowns.

**(C)** Schematic representation of *C. elegans* (note that the membrane of intestinal cells is decorated in green representing a membrane-bound prenylated GFP) and a confocal image of the actual GFP signal in intestinal cells of *C. elegans*. The white square represents the FRAP region.

**(D-E)** Curves and $T_{half}$ values of the normalized fluorescence during a FRAP experiment of worms grown in PA 2 mM plates.

**(F-G)** Quantification of *C. elegans* larval stages and adulthood after 72 h growing at 20°C in F, and tail tip morphology of 1-day adult worms grown on plates with PA-loaded OP50 bacteria in G (n=50 worms/genotype). Examples of the different tail morphologies are shown in Supplementary Fig.3U.

**(H-I)** Average GP index from several images of WT MEFs treated with vehicle, PA 400 µM ± SEW2871 1 µM (S1PR1 agonist) in H and PA 400 µM ± W146 5 µM (S1PR1 antagonist) in I.

**(J)** Relative expression of S1PRs in HEK293 cells measured by qPCR.

**(K-L)** Average GP index from several images of HEK293 cells treated with vehicle, PA 200 µM ± JTE-013 5 µM (S1PR2 antagonist), ± TY52156 5 µM (S1PR3 antagonist) in K and PA 400 µM ± A971432 1 µM (S1PR5 agonist) in L.

**(M)** Relative expression of S1PRs in U-2 OS cells measured by qPCR.

**(N)** Average GP index from several images of U-2 OS cells treated with vehicle, PA 200 µM ± JTE-013 5 µM (S1PR2 antagonist), ± TY52156 5 µM (S1PR3 antagonist).

**(O-Q)** qPCR showing the efficiencies of the S1PR1, S1PR2, S1PR3 knockdowns.

**(R)** Average GP index from several images of NT, AdipoR2 and S1PR1 siRNA HEK293 cells treated with PA 200 µM ± S1P 1 µM.

Data shows mean ± SEM in D-E, H-I, K-L, N and R and ± SD in A-B, J, M and O-Q. *$p<0.05$, **$p<0.01$, ***$p<0.001$.

Supplementary Fig.6 (related to Fig.5).

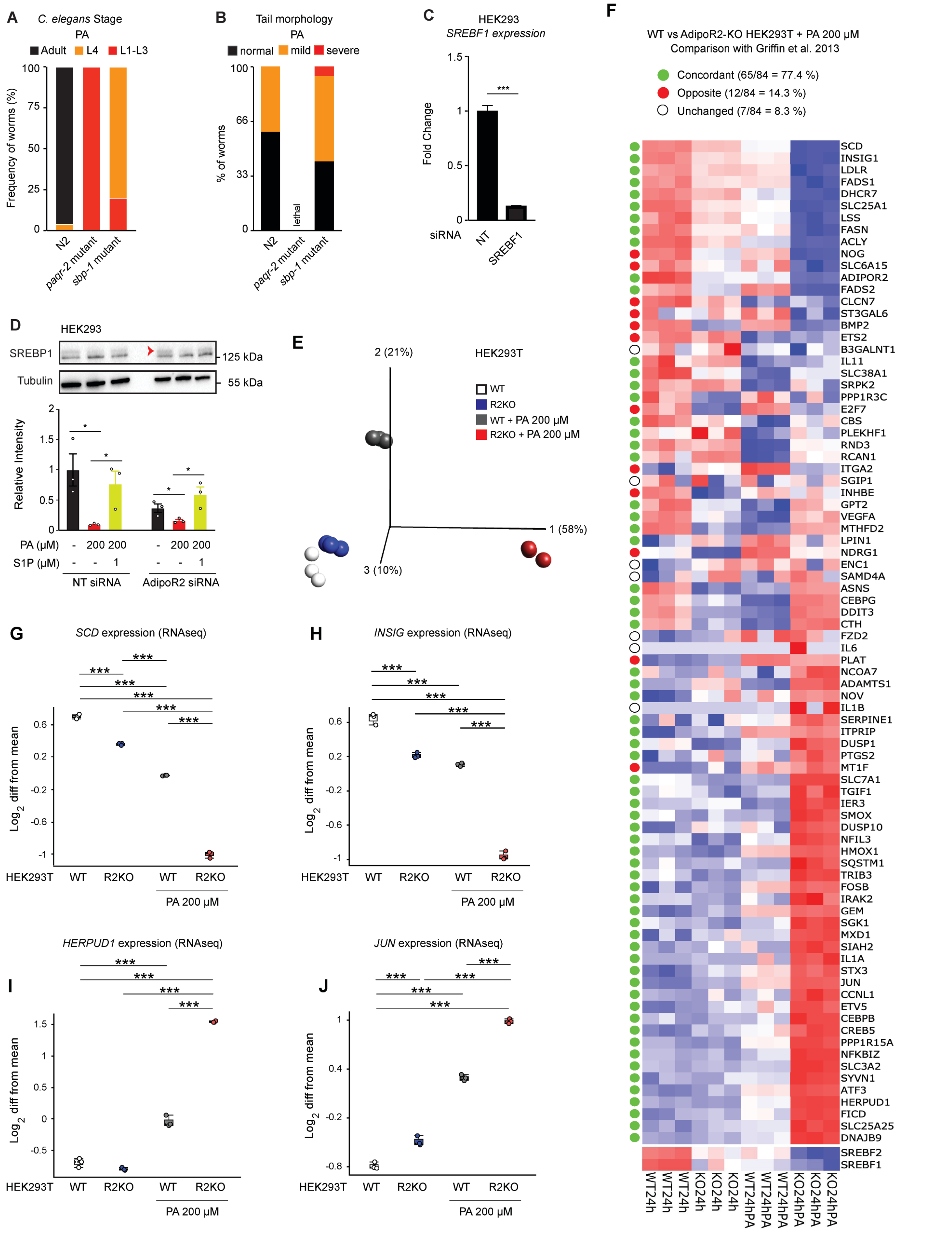

**Supplementary Fig.6. S1P Signaling Via SREBP1 Maintains Membrane Homeostasis in Human Cells and *C. elegans*** (related to Fig.5).
**(A-B)** Quantification of *C. elegans* larval stages and adulthood in N2, *paqr-2* and *sbp-1* mutants after 72 h growing at 20°C in A and Tail tip morphology of 1-day adult worms grown on plates with PA-loaded OP50 bacteria in B (n=50 worms/genotype). Examples of the different tail morphologies are shown in Fig.S3U.
**(C)** qPCR result showing the efficiency of the SREBF1 siRNA knockdown by siRNA in HEK293 cells.
**(D)** Western-Blot and quantification (n=3 experiments) of NT and AdipoR2 siRNA HEK293 cells treated with vehicle and PA 200 µM ± S1P 1 µM. The red arrow points to the precursor SREBP1 band. Note the presence of an unspecific band just below SREBP1. Uncropped blots in Source Data.
**(E)** PCA plot based on 84 SREBP-regulated genes from [1] of WT or AdipoR2-KO HEK293T cells in basal media or basal media supplemented with PA 200 µM for 24 h. Note that genotypes are well separated upon PA treatment.
**(F)** All 84 genes from [1] listed in the order of the PC1, and labelled as to whether their changes in expression matched [1]. *SREBF1/2* were both downregulated in AdipoR2-KO cells and were added at the bottom of the heat map.
**(G-J)** Plots of four genes whose expression its highly altered by the loss of AdipoR2 or SREBP1/2.
Data are represented as mean ± SEM in D and ± SD in C and G-J. *p<0.05, ***p<0.001. See also Supplementary Data 5.

Supplementary Fig.7 (related to Fig.5).

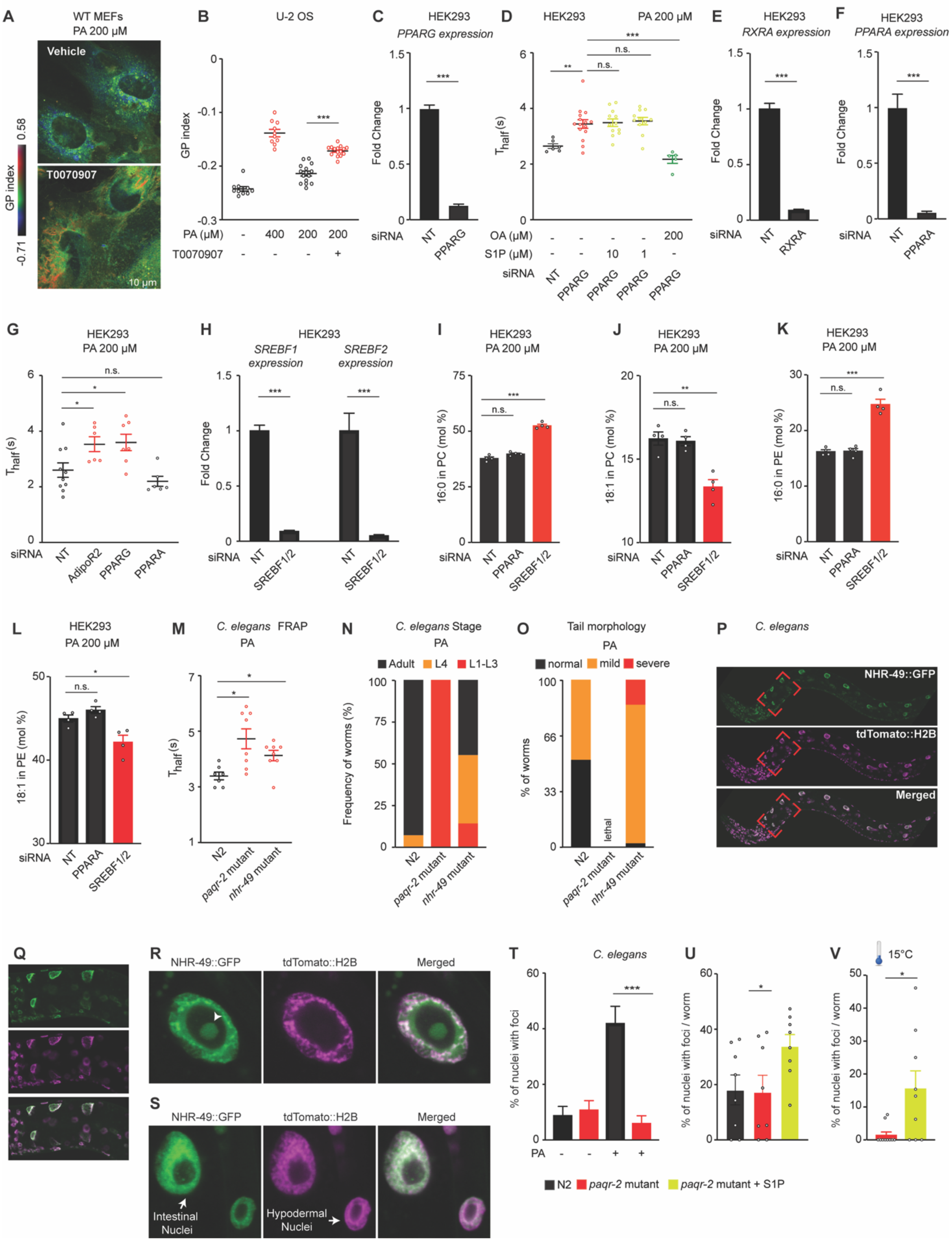

**Supplementary Fig.7. S1P Signaling Via PPARγ Maintains Membrane Homeostasis in Human/Mouse Cells and *C. elegans*** (related to Fig.5).

**(A)** Pseudocolor images of WT MEFs treated with vehicle, PA 200 µM ± T0070907 1 µM (PPARγ antagonist). Quantification of several images is shown in Fig.5D.

**(B)** Average GP index of U-2 OS cells treated with vehicle, PA 400 µM and PA 200 µM ± T0070907 1 µM.

**(C)** qPCR showing the efficiency of the PPARG knockdown.

**(D)** FRAP experiment of NT and PPARG siRNA HEK293 cells treated with PA 200 µM ± S1P (10 or 1 µM) and ± OA 200 µM.

**(E)** qPCR showing the efficiency of the RXRA knockdown.

**(F)** qPCR showing the efficiency of the PPARA knockdown.

**(G)** FRAP experiment in HEK293 cells.

**(H)** qPCR showing the efficiency of the SREBF1+2 knockdown.

**(I-L)** PA 16:0) and OA (18:1) abundance (mol%) in the PC and in the PE of HEK293 cells cultivated as indicated. From Supplementary Data S6.

**(M)** FRAP experiment on N2, *paqr-2* and *nhr-49* mutants fed PA-loaded OP50 bacteria.

**(N-O)** Quantification of *C. elegans* larval stages and adulthood in N2, *paqr-2* and *nhr-49* mutants after 72 h growing at 20°C in N. Tail tip morphology of 1-day adult worms grown on plates with PA-loaded OP50 bacteria in O (n=50 worms/genotype). Examples of the different tail morphologies are shown in Fig.S3U.

**(P-S)** Images of *C. elegans* expressing NHR-49::GFP and tdTomato::H2B (nuclear marker). Panels Q shows zoom-in of P. R shows higher magnification of an intestinal nucleus with nucleolar foci (arrowhead) while S shows intestinal and hypodermal nuclei.

**(T-V)** Quantification of the presence of nucleolar foci in intestinal nuclei of N2 and *paqr-2* mutants grown on control plates (S) (n=248-332 nuclei/genotype) and on plates supplemented with S1P at 20°C (T) and 15°C (V).

Data shows mean ± SEM in B, D, G, I-L, S-U and ± SD in C, E-F, H. *p<0.05, **p<0.01, ***p<0.001. See also Supplementary Data 5-6.

Supplementary Fig.8 (related to Fig.6).

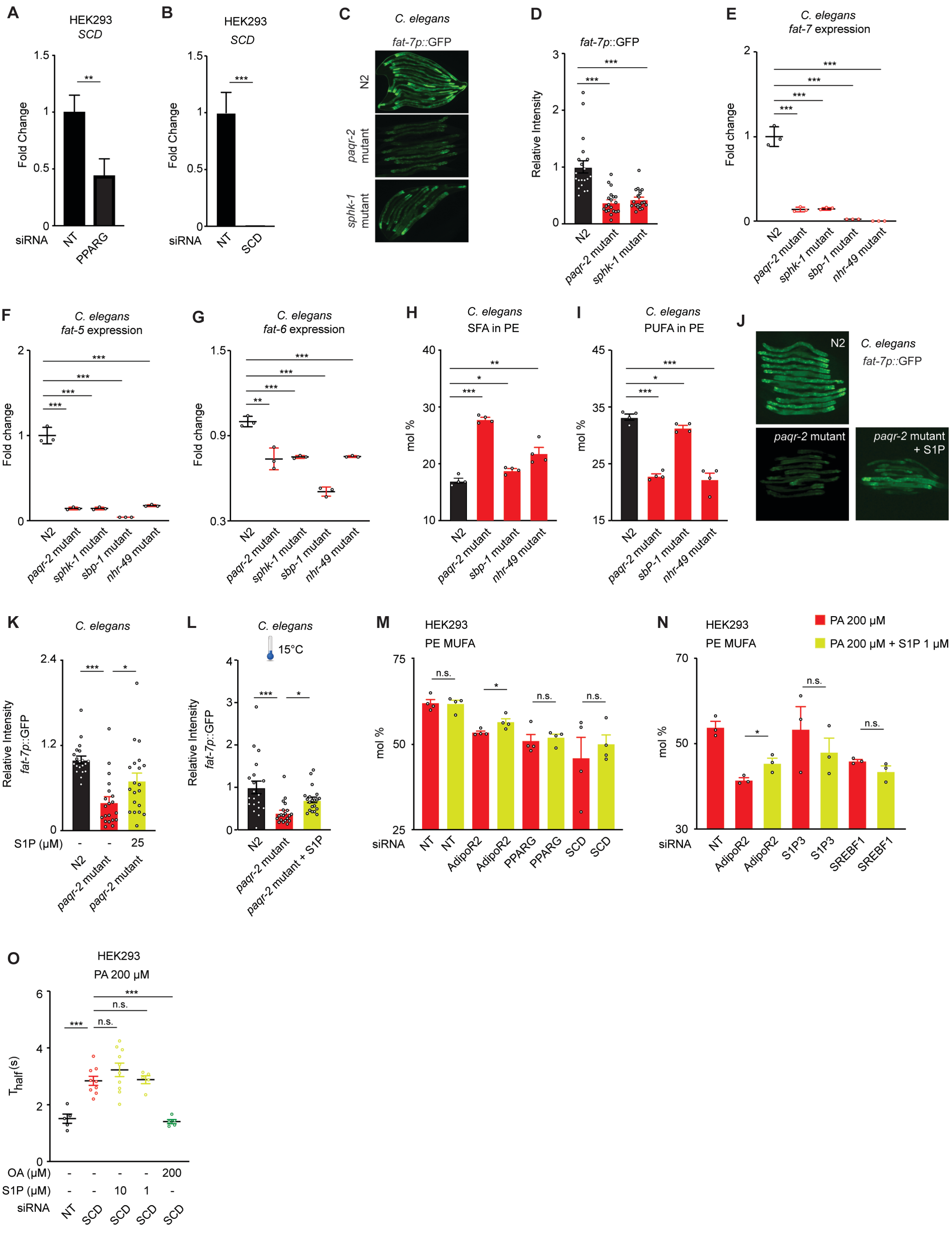

**Figure S8. Role of SCD in Maintenance of Membrane Homeostasis in Human Cells and in *C. elegans*** (related to Fig.6).

**(A)** Relative expression of *SCD* in NT and PPARG sRNA HEK293 cells measured by qPCR.

**(B)** qPCR result showing the efficiency of the SCD knockdown by siRNA in HEK293 cells.

**(C-D)** Representative images of *C. elegans* expressing GFP under the control of the *fat-7* promoter (*fat-7p*::GFP) and quantification of the fluorescence. N2 and *paqr-2* and *sphk-1* mutant worms were grown on control plates for 72 h.

**(E-G)** Relative expression of *fat-7, fat-5 and fat-6* in N2, *paqr-2*, *sphk1* and *nhr-49* mutant worms measured by qPCR.

**(H-I)** PA 16:0) and OA (18:1) abundance (mol%) and in the PE of *C. elegans* grown on PA 2 mM plates. Related lipidomics are shown in Supplementary Data 7.

**(J-K)** Representative images *C. elegans* expressing GFP under the control of the *fat-7* promoter (*fat-7p*::GFP) and the quantification of the fluorescence. N2 and *paqr-2* mutant worms were grown on control plates and plates supplemented with S1P 25 μM for 72 h.

**(L)** Quantification of the fluorescence of *C. elegans* expressing GFP under the control of the *fat-7* promoter (*fat-7p*::GFP) grown at 15°C on control plates and plates supplemented with S1P 25 μM for 144 h.

**(M-N)** MUFA abundance (mol%) and in the PE of HEK293 cells treated with different siRNA and PA 200 μM ± S1P 1 μM. Related lipidomics are shown in Fig.6C-F and in Supplementary Data 6.

**(O)** FRAP experiment of NT and SCD siRNA HEK293 cells treated with PA 200 μM ± S1P (10 or 1 μM) and ± OA 200 μM.

Data are represented as mean ± SEM in D-I, K-O, I and ± SD in A-B. *p<0.05, **p<0.01, ***p<0.001. See also Supplementary Data 5-7.

### SUPPLEMENTARY METHODS

**qPCR** (continuous from main text):
Primers used in the study (some primers were previously described).

| Sequence (5'-->3') | Organism | Purpose - Target | Reference |
|---|---|---|---|
| AGGCAGGGTAAGCTGATTAGCTATG<br>TCCACTGTGTCAGCTTCTCTGTTAC<br>GGGTGGGATTAGATAAATGCCTGCTCT | *Mus musculus* | Genotyping - *Adipor1* | 2 |
| GACGGAGTTTGTATGTGGTAGCGTC<br>TCTCTGCCTTTCCTTTTCATGGCTC<br>GGGCCAGCTCATTCCTCCCACTCAT | *Mus musculus* | Genotyping / *Adipor2* | 2 |
| CTGCCTTTCACCTTGGAGAC<br>CGTTTCCTGGGGATGAGATA | *Mus musculus* | qPCR - *Ddit3* | 3 |
| CATGGTTCTCACTAAAATGAAAGG<br>GCTGGTACAGTAACAACTG | *Mus musculus* | qPCR - *Hspa5* | 3 |
| ATGGCCGGCTATGGATGAT<br>CGAAGTCAAACTCTTTCAGATCCATT | *Mus musculus* | qPCR - *Atf4* | 3 |
| GAGTCCGCAGCAGGTG<br>TAAGACTCCTGCCTGACTGC | *Mus musculus* | qPCR - *sXBP1* | 4 |
| ACTACACAACGGGAGCAACAG<br>GATGGAAAGCAGGAGCAGAG | *Mus musculus* | qPCR - *S1pr1* | 5 |
| CTCACTGCTCAATCCTGTCATC<br>TTCACATTTTCCCTTCAGACC | *Mus musculus* | qPCR - *S1pr2* | 5 |
| TTCCCGACTGCTCTACCATC<br>CCAACAGGCAATGAACACAC | *Mus musculus* | qPCR - *S1pr3* | 5 |
| TGCGGGTGGCTGAGAGTG<br>TAGGATCAGGGCGAAGACC | *Mus musculus* | qPCR - *S1pr4* | 5 |
| CTTAGGACGCCTGGAAACC<br>CCCGCACCTGACAGTAAATC | *Mus musculus* | qPCR - *S1pr5* | 5 |
| GTCTCCTTCGAGCTGTTTGC<br>GCGTGTAAAGTCACCACCCT | *Mus musculus* | qPCR - *PPIA* (housekeeping) | This study |
| TGACCAACAAGGAGATGCGT<br>AATTGTCCGATTTGCTGCGG | *Homo sapiens* | qPCR - *S1PR1* | This study |
| GCCTCTCTACGCCAAGCATT<br>GCAGCCAGCAGACGATAAAG | *Homo sapiens* | qPCR - *S1PR2* | This study |
| TGAAGTCCAGCAGCCGTAAG<br>AGCCAACACGATGAACCACT | *Homo sapiens* | qPCR - *S1PR3* | This study |
| CATCATGGGCCTCTATGGGG<br>AGAGGTTGGAGCCAAAGACG | *Homo sapiens* | qPCR - *S1PR4* | This study |

| | | | |
|---|---|---|---|
| CTTTGTGGCATGTTGGGGC<br>GCGTGTAGATGATGGGGTTCA | *Homo sapiens* | qPCR - *S1PR5* | This study |
| GACTGCCTGATTGACAAGCG<br>ATTCTCGTTCCGGTCCTTGC | *Homo sapiens* | qPCR - *RXRA* | This study |
| ATGCTGGCTATGAGCAGGTC<br>CCCAGACGCCGATACTTCTC | *Homo sapiens* | qPCR - *Sphk1* | This study |
| CCCCGGTTGCTTCTATTGGT<br>GATCTAGGAGCCCGTTCAGC | *Homo sapiens* | qPCR - *Sphk2* | This study |
| GACCAAAGCAAAGGCGAGGG<br>CAGCCCTGAAAGATGCGGATG | *Homo sapiens* | qPCR - *PPARG* | This study |
| GCGAACGATTCGACTCAAGC<br>TCGTCCAAAACGAATCGCGT | *Homo sapiens* | qPCR - *PPARA* | This study |
| GTCTCCTTTGAGCTGTTTGCAG<br>GGACAAGATGCCAGGACCC | *Homo sapiens* | qPCR - PPIA (housekeeping) | 6 |
| GACCTCGCAGATCCAGCAG<br>ATAGGCAGCTTCTCCGCATC | *Homo sapiens* | qPCR - *SREBF1* | 7 |
| GTGCTGTTCCTGACTCCCTG<br>CAGCCTTCTTCTTGGCCTGA | *Homo sapiens* | qPCR - *SREBF2* | 7 |
| TTCGTTGCCACTTTCTTGCG<br>TGGTGGTAGTTGTGGAAGCC | *Homo sapiens* | qPCR - *SCD* | 6 |
| CCATCTGCTTGGTTTCGTGC<br>AGACGGTGTGAAAGAGCCAG | *Homo sapiens* | qPCR - *AdipoR1* | 6 |
| TCATCTGTGTGCTGGGCATT<br>CTATCTGCCCTATGGTGGCG | *Homo sapiens* | qPCR - *AdipoR2* | 6 |
| TCTCGCAGGTTGTGTCTTCC<br>AGCCTCATGGTAAGCCTTGT | *C. elegans* | qPCR - *tba-1*(housekeeping) | This study |
| CAGTTGGATGGGTATTCCTCCT<br>TCCATGAGAGGGTGGCTTTG | *C. elegans* | qPCR - *fat-5* | This study |
| GACCCAGTTCTCGTCTTCCA<br>ATCCGAAATAGTGAGCAGCG | *C. elegans* | qPCR - *fat-6* | This study |
| GCCGTCTTCTCATTTGCTCTC<br>ACGATGATCACGAGCCCATT | *C. elegans* | qPCR - *fat-7* | This study |

#### *C. elegans* Strains

*C. elegans* strains were cultured as in [8]. The wild-type *C. elegans* reference strain N2, *paqr-2(tm3410)*, *sphk-1(ok1097)*, *nhr-49(gk405)* and *sbp-1(ep79)* and the transgene carrying strain EG7865 {*oxTi617 [eft-3p::tdTomato::H2B::unc-54 3'UTR + Cbr-unc-119(+)]*} are available from the *C. elegans* Genetics Center (CGC; USA). The *pfat-7::GFP(rtIs30)* carrying strain HA1842 was a kind gift from Amy Walker [9].

The PHX3258 (*nhr-49::GFP*) strain was created by Suny Biotech (Fuzhou City, China) using CRISPR/Cas9 and carries a modified nhr-49 locus where the end of the coding region is fused in-frame with that of GFP. The altered sequence is as follows (underlined sequences are from the endogenous nhr-49, linker sequences are in bold, GFP coding sequences are in regular uppercase, introns and 3'UTR are in lowercase, and the STOP codon is in italics):
TTGAACAGTGAGCAGAATAATCATATGCTC**AGTAAAGGAGAAGAACTTTTCACTGG**AGTTGTCCCA
ATTCTTGTTGAATTAGATGGTGATGTTAATGGGCACAAATTTTCTGTCAGTGGAGAGGGTGAAGGT
GATGCAACATACGGAAAACTTACCCTTAAATTTATTTGCACTACTGGAAAACTACCTGTTCCATGGgta
agtttaaacatatatatactaactaaccctgattatttaaattttcagCCAACACTTGTCACTACTTTCTgTTATGGTGTT
CAATGCTTcTCgAGATACCCAGATCATATGAAACgGCATGACTTTTTCAAGAGTGCCATGCCCGAAGG
TTATGTACAGGAAAGAACTATATTTTTCAAAGATGACGGGAACTACAAGACACgtaagtttaaacagttcg
gtactaactaaccatacatatttaaattttcagGTGCTGAAGTCAAGTTTGAAGGTGATACCCTTGTTAATAGAA
TCGAGTTAAAAGGTATTGATTTTAAAGAAGATGGAAACATTCTTGGACACAAATTGGAATACAACTA
TAACTCACACAATGTATACATCATGGCAGACAAACAAAAGAATGGAATCAAAGTTgtaagtttaaacatg
attttactaactaactaatctgatttaaattttcagAACTTCAAAATTAGACACAACATTGAAGATGGAAGCGTTC
AACTAGCAGACCATTATCAACAAAATACTCCAATTGGCGATGGCCCTGTCCTTTTACCAGACAACCAT
TACCTGTCCACACAATCTGCCCTTTCGAAAGATCCCAACGAAAAGAGAGACCACATGGTCCTTCTTG
AGTTTGTAACAGCTGCTGGGATTACACATGGCATGGATGAACTATACAAA*TAA*taattccattttttctccca
aaactcttcacctcat. The PHX3258 strain is readily available from the authors and will be submitted to the Caenorhabditis Genetics Center (CGC)

#### *C. elegans* culture conditions

Unless otherwise stated, experiments were performed at 20°C, using the *E.coli* strain OP50 as food source, which was maintained on LB plates kept at 4°C (re-streaked every 6-8 weeks), and single colonies were picked for overnight cultivation at 37°C in LB medium before being used to seed NGM (nematode growth media) plates [10]. Plates containing glucose or S1P were prepared by adding the respective stock solutions to cooled NGM after autoclaving.

#### FRAP in *C. elegans*

FRAP experiments in *C. elegans* were carried out using a membrane-associated prenylated GFP reporter expressed in intestinal cells and using a Zeiss LSM700 inv laser scanning confocal microscope with a 40X water immersion objective. Briefly, the GFP-positive membranes were photobleached over a rectangular area (15 x 4 pixels) using 30 iterations of the 488 nm laser with 50% laser power transmission. Images were collected at a 12-bit intensity resolution over 256 x 256 pixels (digital zoom 4X) using a pixel dwell time of 1.58 µs, and were all acquired under identical settings. The recovery of fluorescence was traced for 25 s. Fluorescence recovery and $T_{half}$ were calculated [11].

**Plasmids for *C. elegans* and Injections**

Wild-type PAQR-2::GFP(*pPAQR-2::N-GFP*) construct have been described elsewhere [12]. PAQR-2::GFP(H511A) construct was generated using PCR-based mutagenesis (Q5-site-directed mutagenesis kit, New England Biolabs) with the following primers: 5'- gctcaactctttcacacatttg-3' and 5'-cgattggaactgaaaaaataaacg-3'. PAQR-2::GFP(H365A, H511A, H515A) construct was generated from PAQR-2::GFP(*pPAQR-2::N-GFP*) using gibson-assembly cloning kit (NEB) with the following primers: 5'-gctacggttgcatgccattctattgaaatggctaaactgt-3' and 5'-gtggcaaagagttgagccgattggaactgaaaaaataaacgt-3' for amplification of the fragment generating H365A and H511A mutations; 5'-tcggctcaactctttgccacatttgtcgtgctcgccgc-3' and 5'-atggcatgcaaccgtagcaaaaaggaatgacattccaaga-3' for amplification of the fragment generating H365A mutation and the rest of the vector sequence. Constructs were injected into N2 worms at 25 ng/µl together with 40 ng/µl *pRF4,* which carries the dominant *rol-6(su1006)* marker used to identify transgenic worms [13].

***C. elegans* Growth**

For length measurement studies, synchronized L1s were plated onto test plates seeded with *E. coli*, and worms were mounted and photographed 144 h (15°C experiments) or 72 h (all other experiments) later. The length of 20 worms were measured using ImageJ [14]. Alternatively, the stages of the worms (L1, L2-3, L4 or adult, n=50).

**S1P Plates and S1P Injections in *C. elegans***

S1P (Avanti Lipids) was dissolved in butanol:methanol (3:1) and sonicated in a water bath at 55°C. S1P stock solution (1.32 mM) was further stored at -80°C. Plates containing S1P were prepared by adding stock solution of S1P to cooled NGM after autoclaving [15]. For injections, 1 µM S1P was directly injected into the *C. elegans* gonad.

**Quantification of Tail Tip Morphology in *C. elegans***

Quantification of the withered tail tip phenotype was done on synchronous 1 day old adult populations, that is 72 h post L1 (n≥50) [16].

***C. elegans* FAT-7 Expression**

Quantification of fluorescence intensity of *pfat-7::GFP* carrying strains were performed on L4 larvae using ImageJ (n≥20) [16].

**Lipidomics Lipid Analysis**

Ceramides, dihydroceramides, glucosylceramides and lactosylceramides were quantified using UPLC-MS/MS on a QTRAP 5000 (Sciex) in positive ESI mode similar to what has previously been described [17]. For this, a part of the lipid extract was reconstituted in

chloroform:methanol:water [3:6:2; v/v/v] and 5 µl was injected into the UPLC-MS/MS system. Molecular species were separated on a BEH C8 column (2.1 x 100mm with 1.7µm particles) from Waters kept at 60 degrees. The mobile phases were water:acetonitrile [70:30; v/v] with 0.1% formic acid as A-phase and acetonitrile:isopropanol [1:1; v/v] with 0.1% formic acid as B-phase. The separation was done at 400µl/min using a linear gradient from 75 to 100% B-phase over 5 minutes. The mobile phase composition was the kept at 100% B for two minutes and then returned to 75% B for 3 minutes for a total runtime of 10 minutes. Quantification of 27 lipid species was done using an external calibration curve made from 16 reference substances (6 ceramides, 4 dihydroceramides, 3 glucosylceramides and 3 lactosylceramides). For accurate quantification internal standards (C17:0 ceramide, C17:0 glucoslyceramide and C17:0 lactosylceramide) were added during extraction. Instrument parameters were: CUR=20, CAD=7, IS=4500, TEM=350, GS1=60, GS2=60, DP=100, EP=10 and CXP=26. Collision energies and MRM-transitions were optimized for each available reference substance.
Sphingosine (Sph) and sphingosine-1-phosphate (S1P) were analyzed using UPLC-MS/MS on a QTRAP 5000 (Sciex) in positive ESI mode similar to what has previously been described [17]. For this a part of the lipid extract was reconstituted in methanol:acetonitrile:water [2:1:1] with 0.1% formic acid and 5µl was inject and separated using a Kinetex C8 column (2.1 x 100mm with 1.7µm particles) from Phenomenex kept at room temperature. The mobile phases were water:acetonitrile [40:60] with 5mM ammonium formate and 0.1% formic acid as A-phase and acetonitrile with 5mM ammonium formate and 0.1% formic acid as B-phase. The separation was performed at 300µl/min using a linear gradient from 0 to 50% B-phase over 5 minutes. The mobile phase composition was then increased to 100% B and held for two minutes before returning to 40% B for 3 minutes for a total runtime of 10 minutes. Quantification was made using external standard curve made from reference substances. For accurate quantification, internal standards ($D_7$-Sph and $^{13}C_2D_2$-S1P) were added during extraction. Instrument parameters were: CUR=20, CAD=7, IS=4500, TEM=500, GS1=60, GS2=60, DP=100, EP=10 and CXP=26. Collision energies were 20 volts for S1P and 13 volts for Sph. The MRM transition used for Sph and $D_7$-Sph were 300.4>282.4 and 307.4>289.4 respectively. For S1P and $^{13}C_2D_2$-S1P the transitions 380.4>264.4 and 384.4>268.4 were used.

Phospholipids, sphingomyelins and triglycerides were all analyzed using direct infusion (shotgun) mass spectrometry on a QTRAP 5500 mass spectrometer (Sciex) as described previously [18-20]. For this a part of the lipid extract was evaporated and reconstituted in chloroform:methanol [1:2; v/v] with 5mM ammonium acetate. For sphingomyelin, the extract was exposed to alkaline hydrolysis (0.1M KOH in methanol at room temperature for 60 minutes) prior to reconstitution in order to remove mass-interfering phospholipids. Infusion was made using a TriVersa NanoMate interface (Advion Bioscience) working in positive mode using 1.2kV and 0.8 psi of nitrogen gas pressure. Precursors of phosphatidylcholine and sphingomyelin were scanned between 600-850 *m/z* and detected using the phosphocholine fragment at *m/z* 184. The phosphatidylethanolamines were detected using neutral loss of m/z 141. Precursors were scanned between 650-850 *m/z*. Triglycerides were detected using multiple neutral loss scanning [21]. Precursors were detected in the range of 800-1000 by monitoring the loss of several fatty acid fragments (between C14:0 to C22:6). The collision energies were optimized per lipid class and were (in volt): PC=45, PE=30 SM=45, TG=35. To achieve a robust signal with sufficient ion statistics, a minimum of 60 scanning cycles were performed for each lipid class. The analyses were performed at a scan rate of 200 Da/s with a stepsize of 0.1 Da.

The generated raw data files (.wiff files) were processed using the LipidView software and identified lipids were quantified against the signal from the lipid-class specific internal standards, which were added during extraction. The internal standards used were d6-TG 16:0/16:0/16:0, PC 17:0/17:0, PE 17:0/17:0 and SM 12:0.

### SUPPLEMENTARY REFERENCES

1. Griffiths, B. *et al.* Sterol regulatory element binding protein-dependent regulation of lipid synthesis supports cell survival and tumor growth. *Cancer Metab* **1**, 3 (2013).
2. Bjursell, M. *et al.* Opposing effects of adiponectin receptors 1 and 2 on energy metabolism. *Diabetes* **56**, 583-593 (2007).
3. Rutkowski, D.T. *et al.* Adaptation to ER stress is mediated by differential stabilities of pro-survival and pro-apoptotic mRNAs and proteins. *PLoS Biol* **4**, e374 (2006).
4. Gomez, J.A. & Rutkowski, D.T. Experimental reconstitution of chronic ER stress in the liver reveals feedback suppression of BiP mRNA expression. *Elife* **5** (2016).
5. Zhang, L. *et al.* A novel role of sphingosine 1-phosphate receptor S1pr1 in mouse thrombopoiesis. *J Exp Med* **209**, 2165-2181 (2012).
6. Devkota, R. *et al.* The adiponectin receptor AdipoR2 and its Caenorhabditis elegans homolog PAQR-2 prevent membrane rigidification by exogenous saturated fatty acids. *PLoS Genet* **13**, e1007004 (2017).
7. Ruiz, M. *et al.* Extensive transcription mis-regulation and membrane defects in AdipoR2-deficient cells challenged with saturated fatty acids. *Biochim Biophys Acta Mol Cell Biol Lipids* **1866**, 158884 (2021).
8. Brenner, S. The genetics of Caenorhabditis elegans. *Genetics* **77**, 71-94 (1974).
9. Walker, A.K. *et al.* A conserved SREBP-1/phosphatidylcholine feedback circuit regulates lipogenesis in metazoans. *Cell* **147**, 840-852 (2011).
10. Sulston, J. & Hodgkin, J. The nematode Caenorhabditis elegan. (ed. W.B. Wood) 507-606 (Cold Spring Harbor Laboratory Press, 1988).
11. Devkota, R. & Pilon, M. FRAP: A Powerful Method to Evaluate Membrane Fluidity in Caenorhabditis elegans *Bio-protocols* **8** (2018).
12. Svensson, E. *et al.* The adiponectin receptor homologs in C. elegans promote energy utilization and homeostasis. *PLoS One* **6**, e21343 (2011).
13. Mello, C.C., Kramer, J.M., Stinchcomb, D. & Ambros, V. Efficient gene transfer in C.elegans: extrachromosomal maintenance and integration of transforming sequences. *EMBO J* **10**, 3959-3970 (1991).
14. Schneider, C.A., Rasband, W.S. & Eliceiri, K.W. NIH Image to ImageJ: 25 years of image analysis. *Nat Methods* **9**, 671-675 (2012).

15. Lee, K. *et al.* In the Model Host Caenorhabditis elegans, Sphingosine-1-Phosphate-Mediated Signaling Increases Immunity toward Human Opportunistic Bacteria. *Int J Mol Sci* **21** (2020).
16. Svensk, E. *et al.* PAQR-2 regulates fatty acid desaturation during cold adaptation in C. elegans. *PLoS Genet* **9**, e1003801 (2013).
17. Amrutkar, M. *et al.* Protein kinase STK25 regulates hepatic lipid partitioning and progression of liver steatosis and NASH. *FASEB J* **29**, 1564-1576 (2015).
18. Jung, H.R. *et al.* High throughput quantitative molecular lipidomics. *Biochimica et Biophysica Acta (BBA) - Molecular and Cell Biology of Lipids* **1811**, 925-934 (2011).
19. Stahlman, M. *et al.* Dyslipidemia, but not hyperglycemia and insulin resistance, is associated with marked alterations in the HDL lipidome in type 2 diabetic subjects in the DIWA cohort: impact on small HDL particles. *Biochim Biophys Acta* **1831**, 1609-1617 (2013).
20. Ejsing, C.S. *et al.* Global analysis of the yeast lipidome by quantitative shotgun mass spectrometry. *Proceedings of the National Academy of Sciences* **106**, 2136-2141 (2009).
21. Murphy, R.C. *et al.* Detection of the abundance of diacylglycerol and triacylglycerol molecular species in cells using neutral loss mass spectrometry. *Analytical Biochemistry* **366**, 59-70 (2007).
